# Variability and stability of large-scale cortical oscillation patterns

**DOI:** 10.1101/093005

**Authors:** Roy Cox, Anna C Schapiro, Robert Stickgold

## Abstract

Individual differences in brain organization exist at many spatial and temporal scales, contributing to the substantial heterogeneity underlying human thought and behavior. Oscillatory neural activity is crucial for these behaviors, but how such rhythms are expressed across the cortex within and across individuals has not been thoroughly characterized. Combining electroencephalography (EEG) with representational similarity and multivariate classification techniques, we provide a systematic characterization of brain-wide activity across frequency bands and oscillatory features during rest and task performance. Results indicate that oscillatory profiles exhibit sizable group-level correspondences, indicating the presence of common templates of oscillatory organization. At the same time, we observed well-defined subject-specific network profiles that were discernible above and beyond the structure shared across individuals. These individualized patterns were sufficiently stable over time to allow successful classification of individuals several months later. Finally, our findings indicate that the network structure of rhythmic activity varies considerably across distinct oscillatory frequencies and features, suggesting the existence of multiple, parallel information processing streams embedded in distributed electrophysiological activity. Together, these findings affirm the richness of spatiotemporal EEG signals and emphasize the utility of multivariate network analyses for understanding the role of brain oscillations in physiology and behavior.

## Introduction

While human brains are very similar, every brain is also distinct. Differences in synaptic strengths and network wiring provide a biological substrate for every individual's unique constellation of memories, beliefs, and personality traits. Magnetic resonance imaging (MRI) techniques have demonstrated not only individual variability of anatomical white matter connectivity (Bürgel et al. 2006), but also marked differences in patterns of correlated hemodynamic activity across distributed brain regions that relate to cognitive functioning (Mueller et al. 2013; Finn et al. 2015; Gordon et al. 2015). In recent years, network approaches applied to MRI data have yielded important insights concerning the brain's macroscopic connectivity pattern, or connectome, and its relation to behavior (Craddock et al. 2013; Sporns 2014). However, a major drawback of MRI is its inherently poor time resolution. In contrast, electroencephalographic (EEG) techniques are sensitive to rapid, millisecond fluctuations in the electromagnetic fields generated by neuronal populations, and are therefore more suitable to examine the highly dynamic nature of rhythmic brain activity. Moreover, high-density EEG combined with spatial filtering techniques offers a reasonable degree of topographical precision, thereby allowing investigation of the “oscillatory connectome” - the pattern of oscillatory interactions across every pair of electrodes. Yet, little is known about the detailed properties of such networks, their variability from person to person, or their longterm stability.

Brain oscillations play a critical role in neural computation and information transfer (Buzsaki 2004), and are causally related to behavioral performance across numerous cognitive domains (Lopes da Silva 2013). Distinct brain rhythms are expressed differently across the brain (Keitel and Gross 2016), and particular frequencies are consistently related to specific functions, such as memory, attention, or decision making (Siegel et al. 2012). Moreover, different aspects of rhythmic activity capture distinct aspects of brain organization. Whereas oscillatory power reflects the degree of local rhythmic activity in a particular frequency band, functional connectivity assesses temporally coordinated activity between brain areas in a band-specific manner. In particular, consistent phase relations between brain circuits are thought to mediate efficient neural transmission on a fast, millisecond timescale (Fries 2005; Fell and Axmacher 2011), while coordinated fluctuations of signal amplitude capture slower aspects of neural communication (Bruns et al. 2000) and relate to the correlation structure observed with functional MRI (Hipp and Siegel 2015). These findings, along with phenomena of cross-frequency coupling (Aru et al. 2014), have instilled the notion that macroscopic electrophysiological signals reflect multiplexed activity, comprising a mixture of multiple parallel communication lines operating in different spectral bands and/or using different coding schemes (Watrous et al. 2015). Importantly, such concurrently present signals can serve functionally distinct behavioral roles and could constitute a fundamental computational principle to increase coding capacity (Schyns et al. 2011; Watrous et al. 2013).

Several studies have begun to investigate rhythmic activity and connectivity from a network perspective. Such investigations have highlighted both rapid changes (Betzel et al. 2012) and the consistency (Chu et al. 2012) of spatially organized oscillatory activity. Moreover, large-scale oscillatory patterns, in line with the multiplexing hypothesis, vary with frequency band, cortical region, and the precise oscillatory characteristic under consideration (Hipp et al. 2012; Brookes et al. 2014; Arnulfo et al. 2015; Siems et al. 2016), as well as cognitive state (Palva et al. 2010; Honkanen et al. 2015). Combined with the possibility of individual differences in oscillatory organization, these findings underscore the brain's enormous complexity. However, how these different sets of observations interrelate is presently not well understood.

Using high-density EEG, we set out to comprehensively characterize the brain-wide structure of multivariate oscillatory networks across all aforementioned dimensions. Adopting representational similarity (Kriegeskorte 2008) and machine learning techniques, we compared individuals’ oscillatory networks based on power, phase-, and amplitude-connectivity, in multiple frequency bands, and during periods of both rest and memory encoding. Our results indicate the existence of highly distinct oscillatory profiles operating in parallel, both within and across individuals. Nevertheless, by assessing network similarity across test sessions spaced hours to months apart, we found oscillatory patterns to be sufficiently stable and unique to allow for successful long-term identification of individual subjects, demonstrating that oscillatory profiles may serve as neural fingerprints.

## Materials and Methods

### Participants

Twenty-one healthy volunteers from the Boston area (8 male, 13 female, mean age ± SD: 22.0 ± 3.0 years, range: 18-31) completed the first visit of this study. Of these, fourteen returned for a follow-up visit several months later (mean: 154 days, range: 109-231). All reported no history of neurological, psychiatric or sleep disorders. Participants were instructed to refrain from consuming recreational drugs or alcohol in the 48 h prior to the study, and to not consume more than one caffeinated beverage on the day of the study. Subjects were compensated monetarily for their participation. All subjects provided informed consent, and this study was approved by the institutional review board of Beth Israel Deaconess Medical Center.

### Protocol

See Figure 1 for an overview of the protocol. The first visit lasted approximately 5.5 h. Subjects reported to the lab at 1 PM, provided informed consent, and were prepared for EEG monitoring. Seated approximately 60 cm from a 27 inch computer display, they underwent a series of rest, memory encoding, and memory retrieval blocks, lasting from 2 to 3 PM. We refer to this first series of rest and task activities as Session A. Next, subjects stayed in the lab and watched a 2 h documentary. Then, from approximately 5.30 to 6 PM, subjects underwent a second series of recordings (Session B), during which they engaged in several additional blocks of rest activity and performed delayed memory tests for the material encoded during Session A.

**Figure 1.**
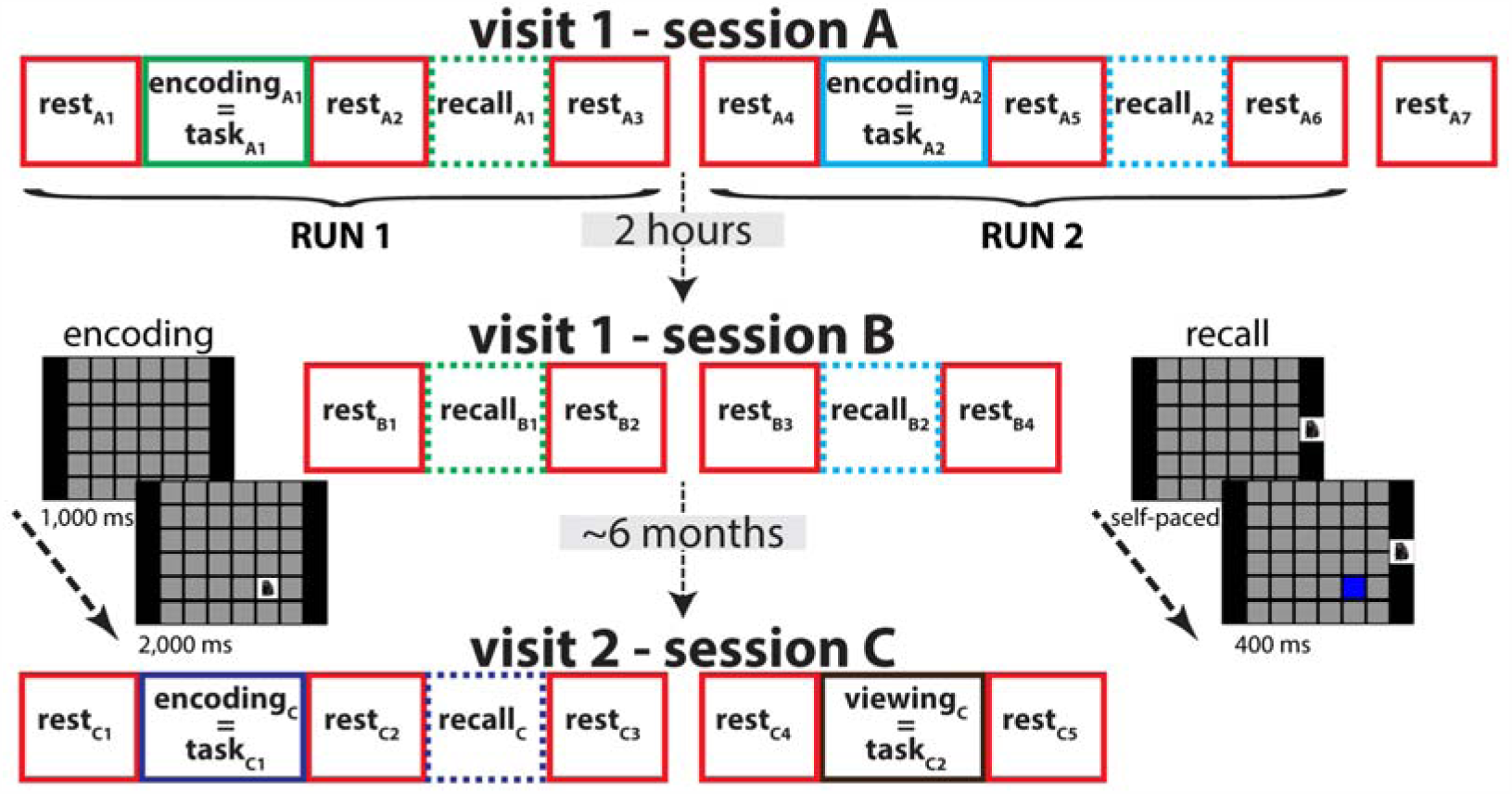
Protocol overview. Sessions A and B were separated by 2 hours, while Session C took place after approximately 6 months. In Session A, there were encoding and recall blocks interspersed with rest periods. In Session B, additional recall blocks were interspersed with rest. In Session C, subjects completed an additional memory task as well as a viewing control task with no memory component. EEG from rest and task blocks (solid lines) but not recall blocks (dashed) was analyzed. During encoding, 36 stimuli were presented, one at a time, on a unique grid location. During retrieval, subjects were cued by presentation of a learned stimulus on the right of the screen, and attempted to select the corresponding location.

After filling out an exit questionnaire, subjects left the lab around 6.30 PM. Participants returning for the follow-up visit several months later (Session C) arrived at the lab at variable times. Following the informed consent procedure and EEG setup, they underwent a series of rest, encoding, retrieval and control blocks for about 45 min. In addition, subjects carried out a 30 min protocol unrelated to the current study. Total duration of the second visit was about 2.5 h.

We use the term “block” to refer to a demarcated period of time associated with a particular behavioral state (i.e., resting, encoding, retrieving, and viewing control blocks; see Rest and Task Details). Importantly, while EEG was recorded during all blocks, retrieval EEG was of very poor quality due to the subjects’ constant movements when operating the mouse. Therefore, and because these data segments were of much shorter duration (~1 min) than the other blocks, we decided not to analyze the retrieval EEG. We adopt the term “data segment” (or just “segment”) to refer to the EEG of rest, encoding, and control (but not retrieval) blocks. Finally, “task” segments refer to data segments from both encoding and control blocks. Note, however, that a control block was only included in Session C, and therefore Session A task segments are equivalent to encoding segments.

For Sessions A and B, each subject's sequence of blocks was organized around the encoding and retrieval of visuospatial associations of two distinct stimulus sets. One set consisted of pictures of animals, the other of vehicles. During Session A, the first five blocks were organized as rest-encoding-rest-retrieval-rest, and pertained to the first stimulus category. Then, this sequence of blocks was repeated for the second stimulus set. The order of stimulus categories was counterbalanced across subjects (animals first: 11; vehicles first: 10). For reasons unrelated to the present report, onscreen instructions then informed participants that later, during Session B, they would be retested only on the first category they were trained on (see below). Then, a final Session A resting state recording was obtained. Thus, a total of 11 behavioral blocks took place, which, after removal of retrieval blocks, resulted in 7 rest segments (rest_A1_-rest_A7_) and two encoding task segments (task_A1_ and task_A2_) for EEG analyses.

After a 2 h interval, Session B began with a reminder that a retest would only be administered for the first encoding category. Then, three blocks were presented in the order rest-retrieval-rest, where the retrieval block reflected the first (and expected) stimulus category. This sequence was followed by a surprise notification that subjects would now also be tested on the second category, and a sequence of rest-retrieval-rest for that category ensued. Thus, for EEG analyses, Session B yielded 4 rest segments (rest_B1_-rest_B4_). An exit questionnaire probed subjects for their learning and retrieval strategies, the amount of time they spent thinking about the stimuli in different phases of the protocol, and their memory and beliefs concerning the expectancy manipulation.

The test expectancy manipulation was originally included to examine whether retest expectation would affect memory consolidation and, consequently, memory performance during Session B, and whether this might be reflected in the EEG.

However, behavioral results did not provide any evidence for this hypothesis and we did not pursue this notion further. Importantly, this state of affairs does not provide a confounding influence hampering interpretation of our results. Beside the fact that memory was not affected by test expectation and we therefore think it unlikely a neural effect would be present, all our EEG analyses were performed across data segments irrespective of expectation category. Moreover, any effect of expectancy would presumably arise after the manipulation is introduced, and could therefore only affect segments rest_A7_ and rest_B1_-rest_B4_. Taken together, we do not believe this manipulation influences the interpretation of the presented analyses.

Session C consisted of one sequence of blocks organized as rest-encoding-rest-retrieval-rest, similar to Session A, and a sequence rest-control-rest. The order of these sequences was counterbalanced across subjects (7 memory first, 7 control first). The stimulus set for memory included 18 animals and 18 vehicles, some of which had been presented during session A. However, all subjects indicated they did not remember any picture-location pairs from their previous visit 4 to 8 months earlier. In the control condition, the same stimulus was repeatedly presented on all locations such that no unique visuospatial memories could be formed. After removal of the retrieval block, Session C resulted in rest segments rest_C1_-rest_C5_ and task segments task_C1_ and task_C2_, one of which was an encoding segment and the other a control segment.

All blocks were presented using custom software written in Java. Instruction screens occurred throughout the protocol during all sessions. Subjects advanced to the next screen by pressing a keyboard button. During rest segments, subjects were instructed to quietly rest and relax for 5 min with their eyes closed, while remaining awake. An auditory tone at the end of each rest segment indicated subjects could open their eyes again.

For memory encoding, subjects were instructed to memorize the location of pictures on a 6 by 6 grid. During the task, a square grid of 36 grey squares, subtending approximately 5° of visual angle per tile, and 31° in total, was continually present on the screen. Thirty-six pictures were then shown, one at a time on a unique grid tile, for 2,000 ms with an interstimulus interval of 1,000 ms. This procedure was repeated for a total of three learning rounds. Picture-location combinations and presentation order were randomized for each subject and were determined at run-time. An encoding block lasted 5 min and 20 s. The control task employed in Session C used the same basic setup as the encoding protocol. Here, however, the same stimulus was placed on each tile, resulting in a perceptually similar experience without the memory demands.

During cued memory tests each picture was presented on the right side of the screen while the grid was visible. The participant was instructed to use the mouse to point the cursor to the tile associated with that picture, at which point the selected tile turned blue for 400 ms. Then, the next trial began. Subjects did not receive feedback on how they did. Retrieval was self-paced and lasted between 1-2 min.

### Data acquisition and preprocessing

EEG was collected using 62-channel caps with channel positions in accordance with the 10-20 system. Two individual Ag/AgCl cup electrodes were attached to the mastoid processes, two were placed around the eyes for electrooculography, and a reference was positioned on the forehead. Channel Afz was used as the ground. An AURA-LTM64 amplifier and TWin software were used for data acquisition (Grass Technologies). Impedances were kept below 25 kΩ and data were sampled at 400 Hz with hardware high-pass and low-pass filters at 0.1 and 133 Hz, respectively.

All subsequent data processing and analyses were performed in Matlab (the Mathworks, Natick, MA) using custom routines and a combination of several freely available toolboxes, including EEGlab (Delorme and Makeig 2004) and Fieldtrip (Oostenveld et al. 2011). EEG recordings were cut into data segments based on triggers derived from the stimulus software. Following the removal of eye channels, data segments were re-referenced to average mastoids, high-pass filtered at 0.5 Hz and notch filtered around 60 Hz to suppress line noise. Noisy channels were interpolated using a spherical spline algorithm (EEGlab: *pop_interp*) and excessively noisy time fragments were removed, resulting in an average segment length across all 364 segments of all subjects of 291 ± 22 s (typical range: 209-325 s; one outlier of 48 s). Independent component analysis (EEGlab: *runica*) was performed and components reflecting eye movements, eye blinks, muscle activity and other obvious artifacts were removed. Next, we applied a spatial Laplacian filter (Perrin et al. 1989) using the CSD toolbox (Kayser and Tenke 2006). The Laplacian reduces the effects of volume conduction by estimating radial current flow, thereby highlighting local aspects of neural processing and allowing for superior estimates of phase coupling and reduced probabilities of observing artificial coupling between electrodes (Cohen 2015; Tenke and Kayser 2015). By decorrelating activity levels across the scalp this approach thus “sharpens” network profiles, thereby improving chances of uncovering subtle network differences.

### Power and connectivity

For each spatially filtered data segment (rest and task) and electrode we estimated power spectral density using Welch's method with 5 s windows and 50% overlap. Power values were dB transformed according to dB power = 10 × log10(power) and averaged across bins for the theta (3-7 Hz), alpha (8-12 Hz), beta (13-30 Hz) and gamma (32-60 Hz) bands. This yielded, for every subject, data segment and frequency band, a vector *V* of length 60 reflecting all electrodes’ power values. Thus, these vectors reflect the network organization of oscillatory power across the scalp. Of note, the dB transformation yielded vectors containing approximately normally distributed values, which is an important assumption for their later use in Pearson correlations. Additionally, as a more concise statistic, we defined global power as the average power across all entries (i.e., electrodes) in a vector: global power = 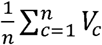, where c is the number of channels (1, 2, …, 60).

For connectivity, we band-pass filtered the segments using the abovementioned cut off frequencies (EEGlab: *pop_eegfiltnew).* We selected these values based on the shape of the filters’ frequency responses, taking care that there was minimal overlap between adjacent pass-bands. Next, we applied the Hilbert transform to each filtered segment and determined the resulting signals’ instantaneous phase and amplitude. We then subdivided the phase angle and amplitude time series into ten equally sized smaller data chunks, and calculated connectivity separately for each chunk. We performed the chunking step to reduce the effect of potential outliers on connectivity estimates, and, for phase synchrony, to allow for non-stationary phase differences within each segment. Chunk length varied across subjects and data segments because data segments had different amounts of artifact removed. While the number of samples affects the signal-to-noise ratio of resulting connectivity estimates, individual chunks were sufficiently long (at least 20 s, except for one outlier with chunk lengths of 5 s) that this variation is unlikely to have had a significant influence on our results.

Amplitude correlations, ranging between −1 and 1, were determined using the Spearman correlation, and were assessed between each channel pair's Hilbert-amplitudes, yielding a matrix *M_amp_* of 60×60 connectivity values for every chunk. We used a nonparametric correlation metric because amplitude envelopes are generally not normally distributed. Phase synchrony was assessed by first calculating the phase difference for every channel pair (j, k) at each sample. We then determined phase synchrony for each channel pair as the length of the average phase difference vector across samples, expressed in the complex plane as: phase synchrony_j,k_ = 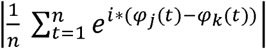 where i is the imaginary operator, *φ* indicates phase (in radians), *t* is the sample, and *j* and *k* index the channels. Phase synchrony values ranged from 0 (random phase relations) to 1 (perfect phase consistency). This resulted in another 60×60 matrix *M_phase_* with phase synchrony values between every channel pair for each data chunk. For both amplitude correlation and phase synchrony, we then averaged connectivity estimates across the ten chunks.

We selected the upper triangles of the symmetrical *M_amp_* and *M_phase_* matrices to count each connection only once. To further limit the effect of spurious coupling on our results we removed from each connectivity matrix the values connecting neighboring electrodes (Fieldtrip: *ft_prepare_neighbours*), removing 192 connections. This yielded, for every subject, data segment, and frequency band, and for both phase synchrony and amplitude correlation, a vector *U* of length 1578 reflecting the degrees of connectivity between all unique channel pairs. Similar to power, we also defined a global connectivity metric as the average across all entries in a vector: global connectivity = 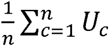, where *c* is the number of connections (1, 2,…, 1578). For topographical plots of connectivity, we averaged the 60×60 matrix *M* (with neighboring connections removed) across one of its dimensions (e.g., row-wise) to obtain the average connectivity between each electrode and all other electrodes.

We observed that connectivity values within each vector *U* were generally not normally distributed. However, the degree of skewness differed depending on frequency band, connectivity metric, data segment and subject. In order to render the data normally distributed for subsequent Pearson correlation analyses, we first added a value of 1 to each connectivity vector entry to ensure all values were positive. We then performed box-cox power transformations to all vectors, where the exponent used for transformation was automatically selected for each vector to minimize the standard deviation of the transformed vector (Box and Cox 1964). Finally, we z-scored the resulting vectors to obtain a standardized appearance when visualized in scatter plots. However, z-scoring does not affect subsequent Pearson correlation statistics. Thus, apart from the power transformation, which was only performed for connectivity vectors, oscillatory power and functional connectivity metrics were processed similarly.

In order to directly compare power and connectivity vectors, which were of different length (*V*: 60, *U:* 1578), we also constructed “power connectivity” vectors of equal size as the connectivity vectors. Specifically, we set the weight of each power “connection” to be the average power of the two involved electrodes. Naturally, this manipulation did not add any novel information, as each of the newly computed values in the larger vector was a linear combination of the original power estimates. As a result, this operation did not influence the similarity among power networks (i.e., power-based network similarity values were identical for vector lengths of 60 and 1578), enabling direct comparisons between power and connectivity metrics.

### Network similarity and statistics

Our basic approach for assessing the similarity of networks involved computing the Pearson correlation coefficient between two vectors. To aggregate network similarities across more than two networks, we used procedures referred to as representational similarity analysis in the modeling and fMRI literature (Kriegeskorte 2008). Specifically, we calculated the Pearson correlation between each unique pair of networks, resulting in a large matrix of similarity scores. Then, for specific questions of interest, similarity values were averaged across the relevant entries and compared to a suitable baseline. Thus, we could compare networks within, between and across subjects, behavioral states, frequency bands, test sessions and oscillation metrics.

In general, we evaluated network similarity statistically on two levels. In “level-1” analyses we determined how much evidence individual subjects showed for a particular phenomenon. Then, these results were summarized across subjects and appropriately tested for significance. In contrast, “level-2” analyses included all subjects’ networks and asked whether there was reliable support for a particular finding at the group level.

For both level-1 and level-2 analyses, we took a data-driven resampling approach to determine whether observed similarity scores across networks were statistically different from what would be expected by chance. For every comparison of interest we constructed a null distribution by repeatedly selecting as many random networks as there were included in the original observation. At each iteration, similarity scores between each pair of selected networks were averaged, thereby creating a surrogate distribution of similarity values. The number of resampling iterations we used depended on the number of unique random samples available. If the number of combinations was over 1,000, a truly random sample was selected for each of 1,000 iterations (i.e., Monte Carlo sampling). When the number of combinations was lower, every unique combination was sampled exactly once (i.e., permutation sampling). Due to the precise mechanics of shuffling, some level-1 analyses could use the same null distribution for every individual, while others required a different baseline distribution for every individual, as outlined below. For every comparison of interest, we defined baseline similarity as the average similarity across permutations (i.e., the center of the null distribution). In case individual subjects used separate null distributions, we further averaged these baselines.

We assessed significance in several ways. First, for some analyses we use a one-sample t-test to compare observed level-1 similarity scores to the average baseline value. This approach tests whether single-subject effects, as a group, are different from the permutation-derived baseline. Second, for both level-1 and level-2 analyses, we used the null distributions to z-score each observation and calculate the associated P values. We then applied the false discovery procedure (Benjamini and Hochberg 1995) to correct for multiple comparisons (i.e., for multiple subjects, frequency bands, oscillation metrics). We used z-based P values, rather than the “raw” permutation-based P values, because the latter often severely underestimated the size of the effect. That is, even with 1,000 iterations the lowest obtainable significance value was P<0.001, while z-based P values provide a more accurate estimate of the distance between an observation and its null distribution. Importantly, for a minority of level-1 analyses, mostly those involving task networks, the number of possible permutations was rather low. As a consequence, the resulting null distributions were based on a limited number of samples and were often not Gaussian-shaped. In these cases, z-based P values did not optimally capture the size of the effect. Moreover, even when observed similarity values were more extreme than the entire null distribution, their raw permutation-based P values were limited by the number of permutations. This state of affairs severely affected subsequent multiple comparison correction with the false discovery procedure. Third, therefore, we additionally present uncorrected, permutation-based P values for these cases in order to offer a complete picture of the pattern of results. In detail, our permutations proceeded as follows for different comparisons, organized by results section:

#### Network consistency within individuals

The consistency of intra-individual network structure for each network type was determined, for rest segments, by selecting a subject's rest networks, computing all pair-wise correlations, and averaging them. For permutation testing, this procedure was repeatedly performed with shuffled subject labels. Then, each subject's observed network similarity was compared to the permutation distribution as explained above. For task segments, the procedure was identical. For rest-task comparisons, we calculated all unique rest-task correlations for each subject. For permutation, subject labels of task segments were kept intact, but rest segment labels were repeatedly shuffled. As a result, a different null-distribution was generated for each subject.

#### Distinct rest and task network profiles across individuals

For each network type, we selected all task segments from all subjects, computed all pair-wise correlations, and averaged them. We did the same for rest, but using only two rest segments from each subject (to equalize the number of rest and task segments). For permutation testing, we repeatedly shuffled the labels of behavioral condition, and recalculated similarity values.

#### Frequency-specific networks for individuals

For every individual, oscillation metric, and behavioral state, we selected all networks of the same frequency, determined each pair-wise correlation, and averaged them. For permutation testing, we repeatedly shuffled frequency labels before recomputing similarity scores. For every individual, observed single-frequency network similarity in different frequency bands could then be compared to a baseline distribution of network similarity scores across frequency bands.

#### Frequency-specific networks across individuals

For every oscillation metric and behavioral state, we selected all networks across all subjects of the same frequency, and determined the average within-frequency network similarity. Frequency labels were then repeatedly shuffled to generate a surrogate distribution of group-level network similarity across frequencies. Observed group-level single-frequency similarity scores were then compared to this null distribution.

#### Distinct power-, phase-, and amplitude-based networks for individuals

For every individual, frequency band, and behavioral state, we selected all networks of the same oscillation metric. For power, we used the “power connectivity” values as explained above. We calculated every pair-wise correlation and averaged them to obtain an estimate of same-metric similarity. We then shuffled oscillation metric labels to generate a null distribution of cross-metric similarity, against which observed vales were compared.

#### Distinct power-, phase-, and amplitude-based networks across individuals

For every frequency band, and behavioral state, we selected all networks of the same oscillation metric across all subjects, and determined the average within-metric network similarity. Oscillatory metric labels were then repeatedly shuffled to generate a surrogate distribution of group-level network similarity across oscillation metrics.

#### Short-term/long-term network stability

For each network type and oscillation metric, we calculated for each subject every unique correlation between segments from the to-be-compared sessions (rest-rest, task-task, and rest-task for Session A-Session B, and Session AB-Session C). For permutation testing, subject labels were kept intact for one session, but were repeatedly shuffled for the other one.

As an additional tool, we employed multidimensional scaling to visualize the similarity of networks, as implemented in Matlab's *mdscale* function.

Multidimensional scaling techniques project high-dimensional data points onto a space of lower dimension while optimally preserving the distances between points (Hout et al. 2013). In our case, each network can be viewed as a point in 60 dimensional (for power topographies), or 1578-dimensional space (for connectivity patterns), and the distance between networks can be expressed as (1 - Pearson correlation), such that more similar networks are closer together. By projecting these points onto two dimensions, relations between networks of different subjects and types that are not apparent from inspecting the full-dimensional data can be approximately visualized. All statistics were done on the full-dimensional data, however.

### Classifiers

We used k-nearest neighbor classifiers (Cover and Hart 1967) as a supervised learning strategy to distinguish between behavioral states within the same session, and to classify subject identity across sessions. In both instances, the algorithm (implemented in Matlab as *fitcknn*) was trained using the correlation distance (1 - Pearson correlation) between each pair of multivariate networks, similar to how we assessed network similarity. Different classifiers were trained for different network types. In the test phase, unseen networks were assigned labels according to the labels in the training set, such that training networks closest to each test network (i.e., more similar networks) contributed more to the final assigned label. This was implemented using an inverse distance-weighting scheme. The number of nearest training networks allowed to vote (i.e., *k*) was set to 5, except for the set of analyses where we investigated classifier performance as a function of *k*.

To classify behavioral states within the same session (i.e., rest vs. task), we used a cross-validated approach in which we repeatedly trained each classifier on the data of all but one subject, and then tested the classifier on the remaining subject's networks. For cross-session subject recognition, we trained classifiers on data from session A and tested them on data from session C. Classifier performance was calculated as the proportion of test networks that were assigned the correct label.

To ascertain whether information pooled across network types improves classification rates of data segments, we combined the information contained by different classifiers. In particular, every individual classifier C_s_ (where s is 1, 2, …., 12 for every single network type: 4 frequency bands × 3 metrics) returns the posterior probabilities that test network X_i_ belongs to training class Y_j_, where j = 1, 2 for behavioral classifiers (rest or task), and j = 1, 2, …, 21 for subject identity classifiers. For every test network X_i_, we averaged the probabilities across classifiers of interest to obtain “class weights” indicating how likely it is network X_i_ belongs to class Y_j_ (note that resulting values do not necessarily sum to 1 and therefore do not reflect true probabilities). Then, the class with the highest class weight determined the assigned class label, similar to how individual classifiers operate.

For subject identification, we further averaged these class weights across the multiple data segments derived from each individual subject. In a final merger step, we combined rest and task networks by taking, for each training class Y_j_, the maximum class weight from the composite rest and composite task classifier, and assigned identity based on the maximum resulting class weight. Classifier performance was evaluated using both binomial tests and permutation-based tests in which we repeatedly shuffled training labels. For various control analyses, we systematically left out network information from particular frequency bands, oscillation metrics, or data segments, prior to merging them.

For searchlight analysis we used the Fieldtrip function *ft_prepare_neighbours* with a neighbor distance of 0.65 to determine the neighborhood structure around each electrode. We then iterated through all electrodes, at each iteration selecting the current electrode, its neighbors, and all non-neighboring connections among them, and trained and tested classifiers on these local networks.

## Results

Twenty-one young, healthy volunteers completed either one or two visits to the lab (Fig. 1). During the first visit, we acquired 60-channel high-density EEG across two sessions (A and B) while subjects underwent a sequence of resting state and memory encoding blocks. During Session A, we recorded seven 5-min segments of quiet, eyes-closed resting activity (rest_A1_-rest_A7_), and two segments of approximately similar duration while subjects encoded arbitrary visuospatial relations in an associative memory task (encoding_A1_ and encoding_A2_). In two corresponding retrieval blocks, subjects’ recall performance was assessed (recall_A1_ and recall_A2_). Following a 2 h break, Session B began, and subjects underwent four additional resting state recordings (rest_B1_-rest_B4_) and performed delayed recall blocks (recall_B1_ and recall_B2_) for the material learned during Session A's encoding_A1_ and encoding_A2_ blocks. Memory performance and its relation to network dynamics are presented in a separate paper (Cox et al., submitted).

After a period of 3.5 to 7.5 months, 14 subjects returned to complete a follow-up Session C, during which five additional resting state (rest_C1_-rest_C5_), and two additional task performance recordings were collected. One of these task segments was another associative encoding task (encoding_c_) to be followed by a corresponding retrieval block (recall_c_), while the second was a passive viewing control task (control_c_) resembling the encoding protocol but without the memory component (and therefore no corresponding retrieval block). We will refer to both encoding and control blocks as general “task” segments (e.g., task_A2_ or task_c1_), except when investigating differences between these block types.

We extracted continuous EEG segments corresponding to the ~5 min blocks of rest and task activity. (We do not investigate stimulus-evoked brain dynamics in this study.) After applying a surface Laplacian filter to data segment to emphasize local neural activity and reduce artificial coupling between electrodes (Cohen 2015; Tenke and Kayser 2015), we determined spectral power at each electrode, and assessed phase synchrony and amplitude envelope correlation between every pair of electrodes. We removed connections between neighboring electrodes for all subsequent analyses to further minimize spurious coupling. We calculated all oscillation metrics (power, amplitude-, and phase-based connectivity) separately for the theta (3-7 Hz), alpha (8-12), beta (13-30) and gamma (32-60) frequency bands. For spectral power, this yielded – for each subject, data segment, and frequency band – a vector of length 60, reflecting all electrodes’ power estimates. For connectivity, the result was a vector of length 1578, reflecting every unique channel pair's connectivity strength. In what follows, we use the term “network” to refer to individual or multiple such vectors, i.e., the set(s) of values reflecting the brain-wide pattern of oscillatory activity across all electrodes or connections. We use the term “network type", to refer to power/connectivity vectors stemming from a particular combination of the oscillatory dimensions under investigation (e.g., amplitude-based beta networks during rest).

### Global and topographical oscillatory activity

In order to obtain an overall picture of oscillatory activity, we first examined global spectral power (i.e., averaged across all channels), for rest and task segments during Session A (Fig. 2A, top). A 4 (frequency) × 2 (rest/task) repeated-measures ANOVA revealed significant main and interaction effects (all P<0.04). In line with typical 1/f frequency scaling, lower frequency bands generally showed greater power than higher bands (paired t-tests: all P_corr_<10^−9^), except for the theta and alpha pair which, due to prominent alpha peaks, showed the reverse pattern during rest [t(20)=- 6.3,P_corr_<10^−5^] and did not differ during task [t(20)=1.7,P_corr_=0.11). Comparing rest and task, theta [t(20)=3.3;P_corr_=0.005] and especially alpha activity [t(20)=8.5;P_corr_<10 ^−6^] were distinctly higher during eyes-closed rest periods compared to task segments, while the reverse was true for gamma power (t(20)=−5.5;P_corr_<10^−4^). Accompanying topographical plots demonstrate the regional contributions to these global effects (Fig. 2A, bottom).

**Figure 2.**
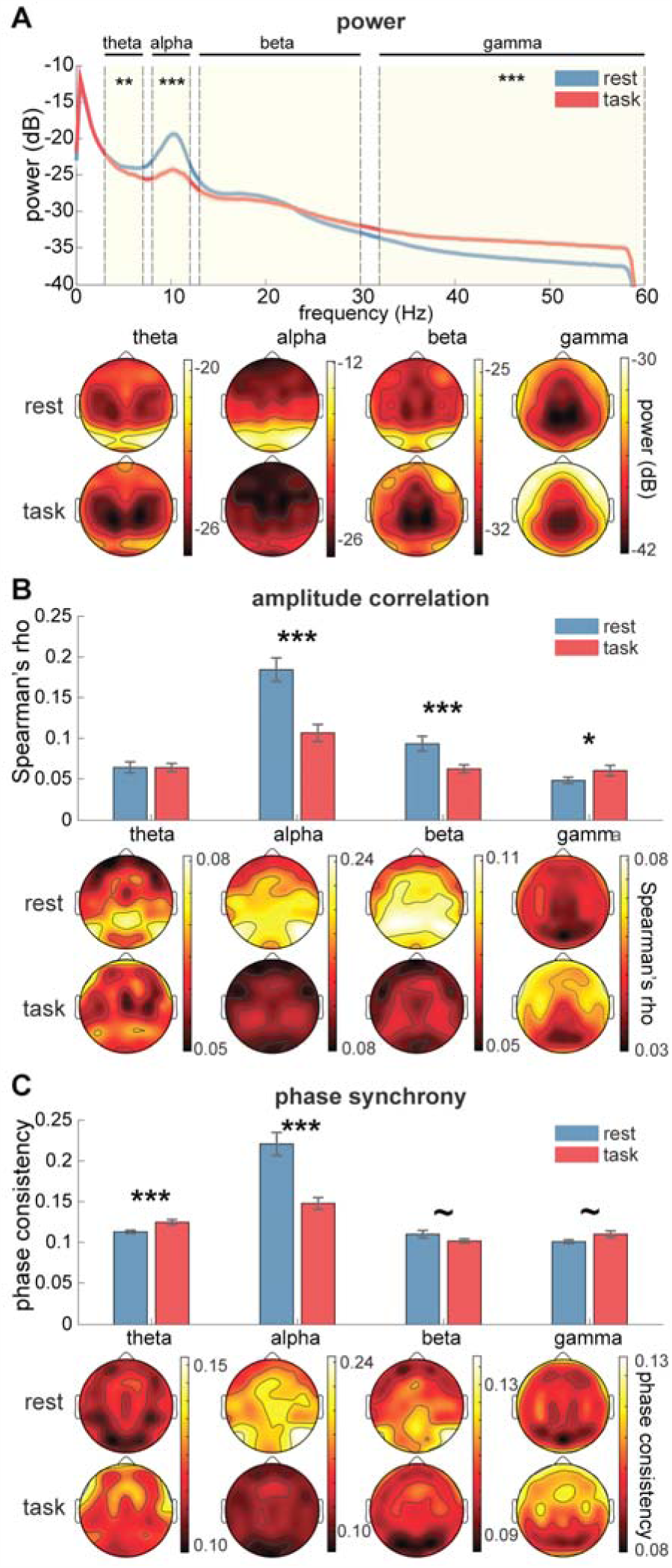
Global and local power and connectivity during session A rest and task segments. **(A)** Top: Power averaged across all electrodes. Rest periods were marked by greater alpha and theta power but reduced gamma activity. Bottom: Topographical distributions of power show typical posterior distributions of alpha activity, with greater power during rest. Global amplitude correlation **(B)**, and global phase synchrony **(C)** averaged across all electrode pairs (top), and as topographical plots where each electrode's connectivity is color-coded according to its average connectivity with all other electrodes (bottom). Symbols indicate significantly (^~^: P<0.1; *: P<0.05; **: P<0.01; ***: P<0.001) different global power or connectivity between rest and task. For details regarding statistical differences between frequency bands see text.

Similarly, to assess overall levels of connectivity, we computed the mean connection strength of each connectivity vector and averaged the resulting values separately for task and resting-state segments. For both amplitude- (Fig. 2B, top) and phase- (Fig. 2C, top) based connectivity metrics, 4x2 ANOVAs yielded reliable main and interaction effects (all P<0.01). Regarding rest-task differences, rest periods exhibited stronger alpha connectivity [amplitude-based: t(20)=6.8,P_corr_<10^−5^; phase-based: t(20)=4.8,P_corr_=0.0004] and beta connectivity [amplitude-based: t(20)=4.7,P_corr_=0.0003; phase-based: t(20)=1.9, P_corr_=0.07]. During task, gamma connectivity was stronger for both metrics [amplitude: t(20)=-2.3,P_corr_=0.04; phase: t(20)=-2.1,P_corr_=0.06], as was phase-based theta connectivity [t(20)=4.0,P_corr_=0.002]. Considering frequencies, all individual frequency pairs demonstrated significant differences for amplitude-based rest connectivity (Fig. 2B, top, blue bars) and phase-based task connectivity (Fig 2C, top, red bars). For amplitude-based task segments and phase-based rest segments, 3/6 and 4/6 frequency comparisons were significant (all P_corr_<0.05). Topographical plots visualizing the average connectivity at every electrode site with the rest of the brain indicate the contributions from specific cortical regions to these global effects (bottoms of Fig. 2BC).

We also examined global and topographical power and connectivity dynamics during Session C and found very similar results. We note that while some features of global connectivity resembled the power profile (e.g., highest values in alpha band during rest), others did not (e.g., greater theta power during rest but stronger theta phase connectivity during task; rest-task differences in beta connectivity but not beta power). To examine in depth if functional connectivity findings could be due to volume conduction we performed several control analyses. However, we did not see a consistent link between global power and global connectivity (Supplementary Text). Analogous to global effects, topographical maps for power, amplitude correlation, and phase synchrony suggest both regional similarities across oscillatory metrics (e.g., posterior activity during rest), and marked differences (e.g., central distributions for beta amplitude-based connectivity not seen for power or phase synchrony).

Combined, findings in this section indicate that global and topographical measures of oscillatory activity vary with frequency and are affected differently by task and rest conditions in separate frequency bands. Moreover, different oscillatory metrics appear sensitive to different aspects of brain dynamics, a notion we will explore more in depth in subsequent sections.

### Similarity of large-scale oscillatory networks

While the previous analyses offer an overall perspective on brain dynamics by examining *absolute* oscillatory activity/connectivity (either topographically or pooled across electrodes/connections), a complementary and more fine-grained approach compares the distribution of oscillatory activity levels across all electrodes or connections between networks. Here, the level of analysis concerns the *relative* distribution of oscillatory activity across the cortex and its consistency from one state to another. In this approach, absolute power/connectivity levels are irrelevant, and not all individual network elements are required to exceed noise levels as signal may still be present in the distributed pattern. Similarly, while we found no evidence for volume conduction affecting connectivity (Supplementary Text), having inflated connectivity estimates due to potential confounds with power does not present a major concern for these network analyses. Again, what matters is the *pattern* of connectivity, not how accurately individual connection weights capture true neuronal synchrony. By then comparing networks across several dimensions (by individual, behavioral state, frequency band, oscillation metric, and across time), new insights concerning the variability and stability of oscillatory organization emerge.

We quantified the degree of similarity between any two networks (i.e., two vectors reflecting brain-wide activity/connectivity patterns) as their Pearson correlation: high similarity between networks indicates a relatively preserved, and therefore stable, configuration of connection strengths or local power across the scalp, irrespective of possible differences in absolute connection strength or power. Illustrative scatterplots for alpha phase synchrony networks demonstrate the generally high correspondence of connection weights between segments derived from the same subject, both within and across behavioral states (Fig. 3AB). In contrast, network similarity between two different subjects was much lower (Fig. 3C). In what follows, we will first characterize oscillatory profiles along all aforementioned dimensions, before turning to the question of how the observed network features may be utilized for subject identification.

**Figure 3.**
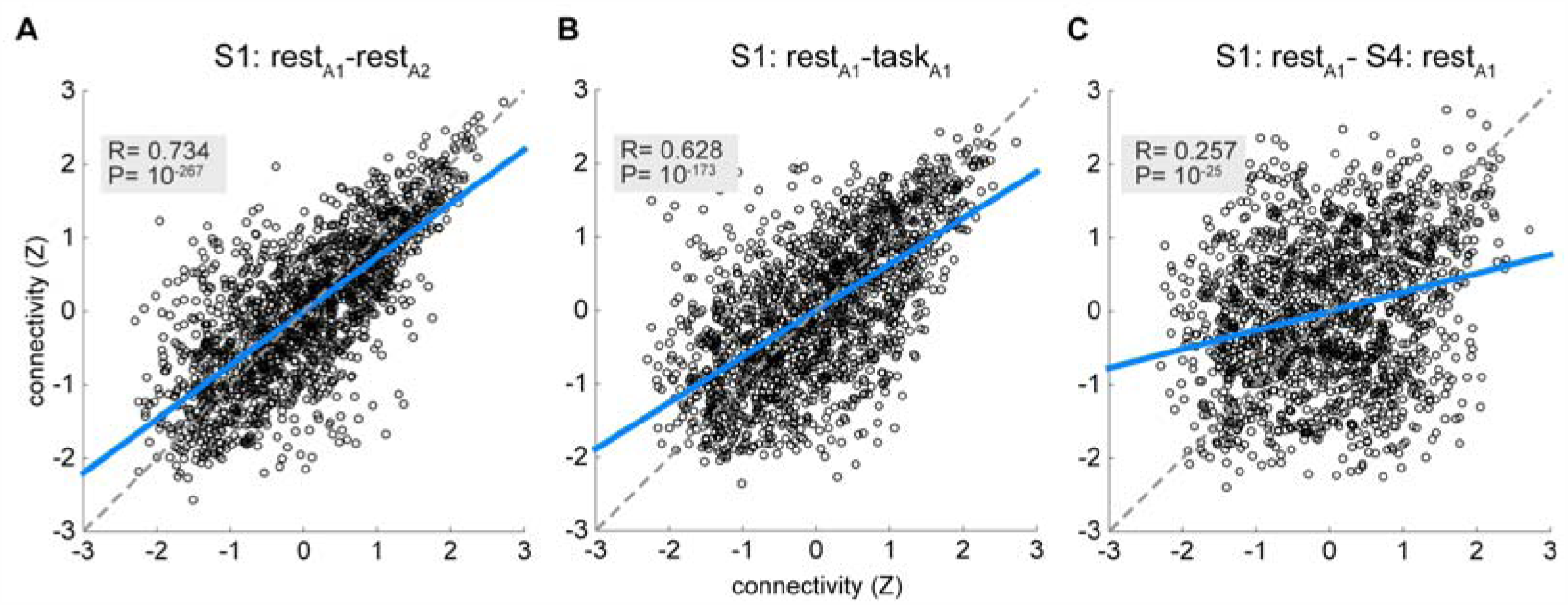
Similarity between alpha phase synchrony networks. Example scatterplots show network similarity between **(A)** a single subject's rest_A1_ and rest_A2_ segments; **(B)** the same subject's rest_A1_ and task_A1_ segments; and **(C)** rest_A1_ from the same subject and the corresponding rest_A1_ of a second subject. Every dot reflects the strength between a pair of electrodes (1578 in total): the Pearson correlation coefficient (R) constitutes the degree of network similarity. Axes indicate z-scored connectivity strength and blue lines reflect least-squares fit. Note that as a result of the large number of data points even modest associations have very low P values.

#### Network consistency within individuals

To examine the notion of intra-individual consistency of network configurations, we assessed the similarity among all Session A data segments obtained from a given subject. For the task conditions, we computed the correlation between task_A1_ and task_A2_ for each subject. For resting states, we computed all unique pair-wise correlations between a subject's 7 (rest_A1_-rest_A7_) segments and averaged the 21 resulting values. We performed this procedure separately for each of 12 network types (4 frequencies × 3 oscillation metrics). Further averaging across subjects, we observed substantial within-subject network similarity, with average Pearson coefficients ranging from 0.49 to 0.98 (Supplementary File 1A; due to the large amount of network comparisons we performed, here, and throughout this report, results are presented at a descriptive level, while detailed network similarity values and statistics are presented in supplementary material). In addition to analyzing networks within behavioral states, we determined intra-subject network consistency between segments of rest and task. Here, we calculated, for every subject, the average correlation between each of the 14 unique pairs of rest-task segments. Compared to similarity of networks from the same behavioral state, correlation values were reduced, but still sizable (range: 0.33-0.73; Supplementary File 1A). Overall, intra-individual similarity scores indicate that network profiles are highly correlated across sequential blocks of time, with strong effect sizes within a behavioral state, and moderate to strong effects between rest and task.

To evaluate whether these correlation scores indicate intra-subject network consistency beyond what is expected by chance, we adopted a resampling approach in which we randomly selected networks from the pool of all subjects. Keeping network type (i.e., frequency, oscillation metric, behavioral state) constant, we repeatedly permuted subject labels to generate a null distribution of similarity values (see Methods). Null distributions for rest-rest and rest-task comparisons are shown in Fig. 4A, C, and D. Individual subjects’ values (orange bars) had far higher similarity values than expected by chance, and were often the most extreme scores. As a complementary tool, we employed multidimensional scaling techniques (see Methods) to visualize the relatedness of these networks (Fig. 4BE). Here, each colored dot reflects a network's position in high-dimensional space, and smaller distances between dots reflect greater similarity between networks. These plots, with each subject coded in a separate color, demonstrate that oscillatory profiles from the same individual are tightly clustered together in multivariate space, indicating that network structure is highly stable for a given individual.

**Figure 4.**
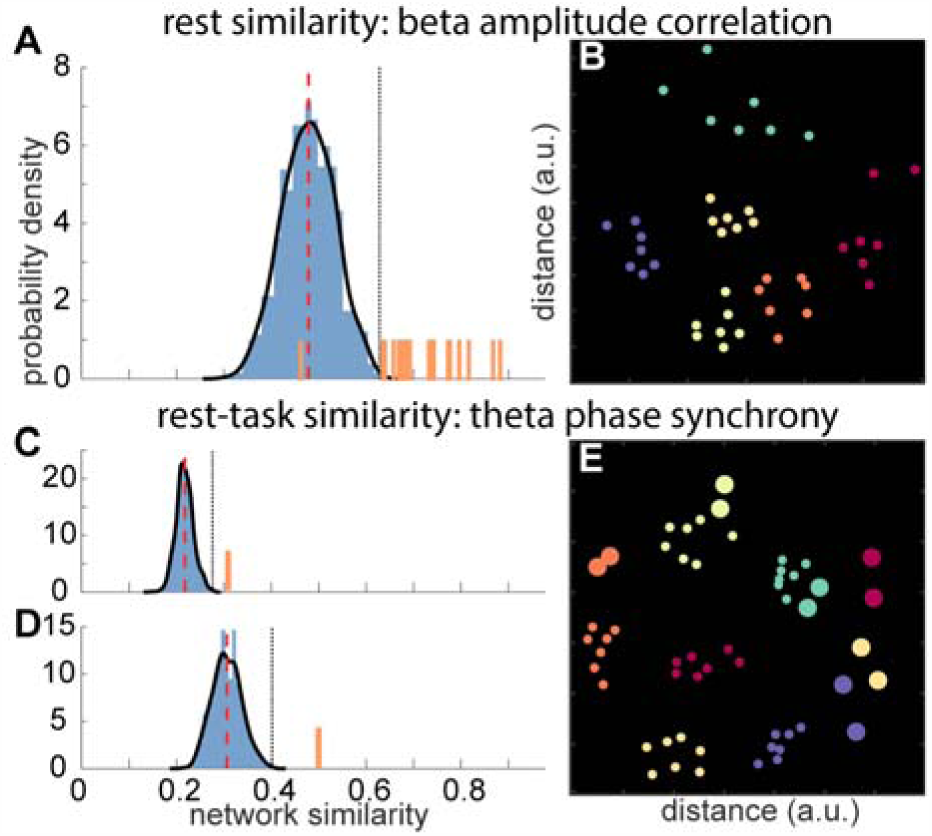
Within-subject similarity of rest and task segments in Session A. Top: Similarity (Pearson's R) of rest segment networks based on amplitude correlation in the beta band. **A:** Observed within-subject similarity values (orange bars) are much higher than for the null distribution generated by resampling across subjects (red line: mean of null distribution; black dotted line: maximum value in distribution). **B:** Multidimensional scaling plot (see Methods) shows similarity between networks for same network type as in **A** as distances between dots. Each color represents a single individual. Dots of the same color are generally clustered together, reflecting high intra-individual network similarity. For visualization purposes only six subjects are plotted, although clustering is equally present when including all 21 subjects. **<u>Bottom:</u>** Similarity between task and rest segments for theta phase synchrony networks. Each subject's similarity score across behavioral states was compared to its own null distribution. Distributions for two subjects **(C** and **D)** show much higher within-subject similarity between rest and task structure (orange bars) than expected by chance. **E:** Distance plot for rest-task similarity, as presented in **C** and **D.** Smaller dots indicate rest networks (as above) and larger dots signify task networks. For several subjects, their 2 task segments are close to their 7 rest segments, indicating a close correspondence between network structures across behavioral states. At the same time, task networks from different subjects tend to cluster together to the right of the plot, suggesting group-level differences between task and rest networks. Again, only 6 subjects are plotted for visualization purposes.

We assessed the significance of this apparent within-subject network stability in two ways. First, to examine group-level effects, we performed a series of one-sample t-tests comparing the distribution of observed similarity scores across subjects to a null-hypothesis baseline defined as the average similarity across permutations. This procedure was performed separately for each network type. For reference, these baseline scores are visualized as red dashed lines in the center of the null distributions presented in Fig. 4 (ACD). For all frequency bands, oscillation metrics, and data segment comparisons (within-rest, within-task and rest vs. task) this yielded highly significant results (all P<0.002, Supplementary File 1A), indicating that Session A power and connectivity profiles are more similar within than between subjects.

Second, we used the null distributions to z-score each participant's network similarity estimates and calculate the associated P values. We then used the false discovery rate (Benjamini and Hochberg 1995) to correct for the multiple tests we performed, thus determining significance for each individual separately. Results indicated 90-100% of individual subjects, depending on network type, displayed above-chance (P_corr_<0.05) network similarity within their resting state recordings for all oscillation metrics and frequency bands. For task segments, within-subject network stability was significant for 55-90% of subjects across network types, except for beta and gamma power profiles. Concerning rest-task similarity, 65-100% of subjects exhibited significant network stability across these behavioral states. We note that for task-task similarity, percentages of subjects reaching significance increased for 9/12 (3 metrics × 4 frequency bands) network types when using more lenient uncorrected thresholds. Details of subject proportions meeting corrected and uncorrected thresholds for each combination of oscillation metric, frequency band and behavioral state are reported in Supplementary File 1A.

We repeated these analyses using only two rest segments, matching the number we had available for task segments, thus removing any potential bias due to different amounts of rest and task data. Our results were very similar (Supplementary File 1B). We also assessed Session B intra-subject resting state similarity (Supplementary File 1C), and Session C intra-subject rest, task, and rest-task similarity (Supplementary File 1D). These analyses showed highly similar and consistent patterns of results, providing independent confirmation of intra-subject network consistency across the same recording session, both within and between behavioral states, and, equivalently, sizable between-subject variability in network organization.

#### Distinct rest and task network profiles across individuals

Analyses of rest and task network patterns showed striking differences between these behavioral states. Upon visual inspection of the distance plot in Fig. 4E, it appears task segments from different subjects tend to cluster together (larger dots in the lower right of the plot), suggesting the existence of group-level differences between task and rest network structure. To investigate this further, we calculated group-level similarity across all subjects’ rest segments, and separately, across all task segments, and compared these values to a baseline distribution of similarity scores obtained through resampling from the combined pool of rest and task segments across subjects. This approach was taken separately for each network type.

Observed similarity scores during Session A varied depending on the oscillation metric, frequency band, and condition analyzed (Supplementary File 2A), but overall, 10 of the 12 (3 metrics × 4 frequency bands) network types exhibited significantly clustered network configurations, during either rest, task or both. These findings indicate that network organization across individuals within a behavioral state (rest, task) is more similar than would be expected by chance. We repeated this procedure for the rest and task segments from Session C and obtained similar results (Supplementary File 2B). Thus, in addition to individual differences in network organization, both rest and task networks share common profiles across subjects.

We asked whether the observed group-level rest-task differences would allow us to predict behavioral state from network structure. For each [oscillation metric/frequency band] combination, we trained a *k*-nearest neighbors classifier (see Methods) on Session A rest and task networks from all subjects. In a cross-validated approach, we repeatedly left out each subject's networks from the training procedure and allowed the classifiers to predict their associated behavioral state. We obtained significantly greater than chance (50%) performance for all 12 network types (binomial tests: all P_corr_<0.04). Recognition rates ranged from 60% for gamma amplitude- and power-based networks, to 88% for alpha phase- and amplitude-based networks (Supplementary File 2A). Average performance across 12 classifiers was 77 ± 10%. Rest networks were more accurately classified than task patterns (83 ± 19% vs. 71 ± 12%), although the difference was not significant [(t(11)=1.85, P=0.09]. Merging evidence from individual classifiers, each based on a different network type (see Methods), we obtained a classification rate of 92%, indicating different network types are sensitive to different aspects of rest-task differences. Repeating these analyses for Session C, we again found considerable evidence for distinct task and rest-based networks (Supplementary File 2B).

In sum, these observations demonstrate that, in addition to subject-specific networks that remain stable from rest to task execution, oscillatory profiles of these behavioral states nonetheless exhibit global task-rest differences that can be discerned at the group level.

#### Frequency-specific networks for individuals

The previous findings demonstrated reliable intra-individual network consistency for all examined frequency bands. However, this leaves unanswered whether an individual's oscillatory networks are similar across frequencies, which would suggest that they derive from the same intrinsic network activity, or whether distinct spectral bands are independently organized, suggesting the existence of multiple parallel modes of neural processing.

We used our resampling approach to address this issue, comparing within subjects the similarity of single-frequency networks to the similarity of networks selected randomly across frequencies. For Session A resting states, every subject showed significantly enhanced network similarity within one or more frequency bands than across bands (Supplementary File 3A), indicating that the involved frequency-specific networks differed reliably from each other. We found this to be the case for all three oscillation metrics. In terms of frequency, networks in the alpha range were most distinctly clustered (for all oscillation metrics), and, in terms of oscillatory feature, networks based on phase synchrony showed most reliable between-frequency differences. Overall, across oscillation metrics and frequency bands, 70-100% of subjects showed significantly greater than chance within-frequency consistency (Supplementary File 3A). To assist interpretation, Fig. 5A uses multidimensional scaling to visualize single-frequency clustering for a sample subject's phase-based rest segments, where different colors indicate different frequencies and shorter distances between dots indicate greater network similarity. Repeating these analyses for rest segments from Sessions B and C, >90% of subjects showed significant single-frequency clustering for one or more frequency bands, with subject percentages across all network types ranging between 50-100% for Session B (Supplementary File 3B), and between 35-90% for Session C (Supplementary File 3C).

**Figure 5.**
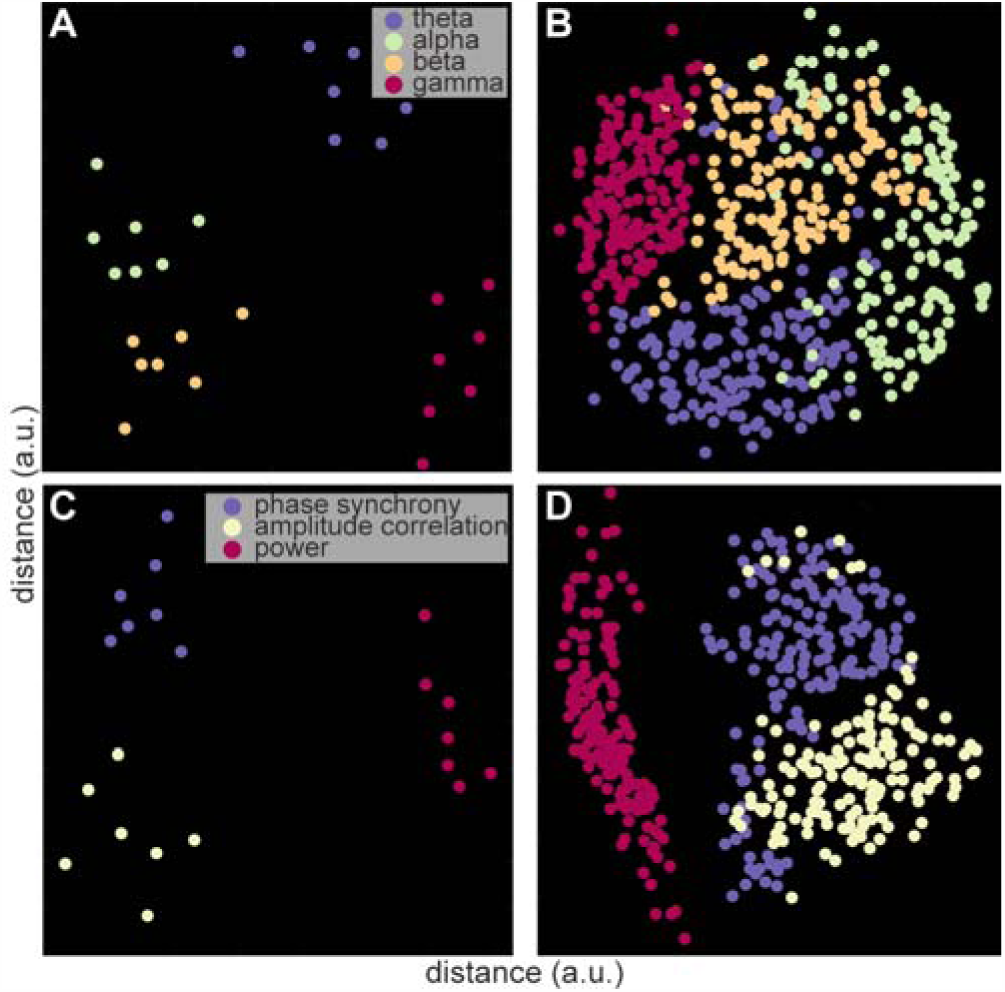
Frequency- and oscillation metric-specific clustering of Session A resting state networks. Single subject-level **(A)** and group-level **(B)** frequency clustering of phase-based networks indicating greater similarity of same-frequency than between-frequency oscillatory profiles. Single subject-level **(C)** and group-level **(D)** oscillation metric clustering in the beta range. Note how power topographies are distinctly different from both connectivity-based metrics.

We performed an analogous set of analyses on each individual's task segments, for both Sessions A and C. The observed same-frequency similarity was often the most extreme score of all possible permutations and, depending on oscillation metric and session, 65-90% of subjects exhibited frequency-specific task networks for at least one frequency (Supplementary File 3A and 3C). All told, these findings strongly indicate that connectivity and power profiles differ across frequencies within individuals, for both rest and task, suggesting large-scale oscillatory activity is organized in frequency-specific manner.

#### Frequency-specific networks across individuals

Next, we turned to the question of whether frequency-band specificity extends to the group level. Given that we observed statistically distinct oscillatory profiles for different individuals, it does not trivially follow that single-frequency networks across subjects are more similar than expected by chance. Using our permutation approach, we observed significantly enhanced network similarity within frequency bands for all oscillation metrics during rest segments from Session A (Supplementary File 3A). Correspondingly strong clustering was visible in multidimensional scaling plots (Fig. 5B). Similar group-level correspondences were found for Session B and C rest segments (Supplementary File 3B and 3C). Task segments from Sessions A and C also demonstrated significant group-level clustering, except for theta and alpha connectivity networks (Supplementary File 3A and 3C). Thus, these findings suggest not only the existence of within-subject, frequency-specific networks, but also the presence of canonical frequency-dependent networks across subjects.

#### Distinct power-, phase-, and amplitude-based networks for individuals

We also see important distinctions among networks based on the oscillation metric employed. As explained in the introduction, estimates of power, amplitude correlation, and phase synchrony should be sensitive to distinct facets of oscillatory activity and communication (Bruns et al. 2000; Cohen 2014; Watrous et al. 2015; Bastos and Schoffelen 2016). However, whether this separation extends to the level of brain-wide EEG patterns is an open question. Using our permutation approach, we asked whether network configurations were reliably more similar when they derived from a single oscillatory metric than when the networks were randomly selected across oscillatory categories.

Overall, single-metric correlation values for Session A resting state networks were greater than the average correlation stemming from permuting across oscillatory metrics (Supplementary File 4A). All subjects displayed significant single-metric clustering for all oscillation metrics for the theta, beta and gamma bands, while for alpha 90, 95 and 100% showed significant similarity for phase-, amplitude-, and power-based networks, respectively. Fig. 5C displays a corresponding distance plot for a single subject's resting beta networks. We repeated these analyses for Sessions B and C rest networks and found similarly high proportions of subjects with significantly distinct network types (Supplementary File 4B and 4C). Thus, these findings demonstrate that oscillatory profiles based on different oscillatory features are reliably distinct, even when derived from the same frequency band, for almost all individuals.

#### Distinct power-, phase-, and amplitude-based networks across individuals

Next, we asked whether networks based on different oscillation metrics are consistently distinct across subjects. Permutation testing demonstrated this be the case for all frequencies except alpha, for both rest and task segments, for all metrics, and during all sessions. For alpha, at least one metric failed to reach significance in each session (Supplementary File 4A-C). Fig. 5D displays the group-level similarity of resting beta networks during Session A across oscillation metrics. These data demonstrate that power-, phase-, and amplitude-based network patterns are differently organized, both within and across individuals.

As seen in Fig. 5CD, power networks differed substantially from connectivity networks in general, with phase- and amplitude showing less difference. We therefore repeated the preceding set of analyses excluding power networks and found that 80-100% of individual subjects showed significant clustering for one or both metrics in the theta, alpha and beta bands, with lower proportions of 30-60% for gamma (Supplementary File 4D-F). Generally, phase synchrony networks showed more reliable within-subject same-metric network consistency than amplitude-based networks. In contrast, group-level network consistency was significant mostly for amplitude-based networks. These effects again occurred for both rest and task networks, and across all sessions and frequency bands. Thus, direct comparisons between functional connectivity networks based on mathematically and theoretically distinct measures of neural communication confirm the distinctiveness of these networks, both across and within individuals.

#### Short-term network stability

The previous analyses demonstrated intra-subject network consistency within a single recording session, but did not address the stability of these patterns across longer periods. We first studied the stability of network structure across sessions by comparing networks from Sessions A and B, which were spaced 2 h apart. We compared observed within-subject, between-session similarity scores to null distributions in which we paired each subject's networks from one session with networks randomly selected across subjects from the other session. We did this for all network types, and separately for rest A-rest B and task A-rest B comparisons.

Both sets of analyses yielded within-subject similarity values that were consistently greater than those obtained by chance (Supplementary File 5A). At the group-level, higher than expected network stability occurred for every network type, and for both rest-rest, and task-rest comparisons (one-sample t-tests: P < 0.005 for each comparison). At the single subject-level, 90-100% of individuals reached significance for rest-rest stability, depending on network type, while 50-100% showed significantly stable network organizations between rest and task segments across the 2 h interval. Interestingly, this pattern of results did not deviate substantially from the within-session analyses (Supplementary File 1A). Across network types, proportions of subjects showing significant same-session versus between-session stability were similar for rest segments [96.4 ± 4.1 vs. 97.6 ± 3.2%, t(11)=-1.4, P=0.19, paired t-test], and decreased over time for rest-task comparisons [86.9 ± 9.3 vs. 72.2 ± 15.0%, t(11)=6.6, P<10^−4^]. These findings suggest a modest change in oscillatory brain patterns over a 2 h period.

#### Long-term network stability

Whether the within-subject stability of oscillatory brain patterns seen across 2 h reflects state or trait differences is unclear. State differences based on any of a number of psychological or physiological parameters (e.g., mood, hours slept the previous night) might form the basis of these individual differences. To examine this further, we used Session C data to analyze stability of individual subjects’ network structure across a period of months. We performed four sets of comparisons between Session C and Sessions A and B: rest-rest, task-task, rest-task, and task-rest.

Because rest segments from Sessions A and B were highly similar, we treated these data as stemming from one session.

Assessing single-subject-level statistics across months, we observed sizable proportions of individual subjects with significant similarity scores. Between 15 and 60% of subjects exhibited significant cross-session network stability from Sessions A and B to Session C for rest-rest and rest-task comparisons, depending on frequency band and oscillation metric (Supplementary File 5B and 5C). Values for task-task and task-rest comparisons were somewhat lower and ranged from 0 to 40%. At more lenient thresholds (P<0.05, uncorrected), the number of subjects showing consistent network stability increased, but for no network type did the proportion rise above 60%. Clearly, subject proportions reaching significance for long-term inter-session stability were reduced compared to same-session consistency. Compared across all oscillation metrics and frequency bands, subject proportions reaching significance decreased from 94.4 ± 4.1% to 37.5 ± 11.4% for rest segments, and from 76.6 ± 10.6% to 18.4 ± 7.8% for task data [t(11)=19.5, P<10^−9^ and t(11)=14.2, P<10^−7^]. Still, for 47 out of the total 48 long-term cross-session comparisons we performed, one or more subjects showed evidence for significant network stability. In sum, while we found substantial evidence for long-term stability of oscillatory networks within individuals, a clear reduction in network similarity was also apparent.

### Identifying individuals from oscillatory patterns

#### Data segment classification

The permutation approach described in the last section asked whether a subject's Session C networks were more similar to his or her own Session A/B networks than to Session A/B networks taken across participants. We next asked whether a supervised learning technique might be more sensitive to long-term network stability by capitalizing on only the most similar networks across sessions. To address this question, we trained a set of *k*-nearest neighbors classifiers, one for each frequency band/oscillation metric/behavioral state combination, on Session A and B network configurations. We then allowed the trained classifiers to predict subject identities for Session C networks. Of note, while data of only two-thirds of the original volunteers was available to assess classification accuracy, each classifier was trained on all 21 subjects’ network patterns and was allowed to predict any of the 21 identities.

Trained on all available session A and B segments (11 from rest, 2 from task), classifier performance for Session C segments (5 rest and 2 task) was significantly above the chance rate of 4.8% (1/21), ranging from 33% for theta amplitude-based networks during rest, to 79% for alpha phase-based patterns during task (Table 1). All classifiers were significant at P<10^−10^ (binomial test), and returned a more extreme result than observed across 1,000 iterations of reshuffling subject labels (all P<0.001). On the whole, the set of rest classifiers performed similarly to the set of task classifiers [t(11)=1.7,P=0.12], For reference, we compared these classification rates to rates obtained with the uncorrected (and most lenient) threshold of the similarity-and-permutation approach. Without exception, classifiers performed better [t(11)=4.5 and t(11)=7.5 for rest and task, both P<0.001]. In a control analysis, we limited the number of training and test segments available to rest classifiers to two of each (from Session A), to match the number of segments used for task classifiers (Supplementary File 6). As expected, this resulted in reduced performance for rest classifiers, but also for rest compared to task classifiers [t(11)=-3.6,P=0.005], although the poorest performing classifier still correctly identified 25% of segments and performed better than all attempts with reshuffled training labels (both binomial and permutation: P<0.001). In sum, these findings demonstrate that oscillatory network patterns carry substantial information for classification across months, for all oscillation metrics and frequency bands, and for periods of both rest and task.

**Table 1.** Classifier performance for all oscillation metrics, frequency bands and behavioral states. Numbers indicate percentage of data segments correctly identified. All classifiers performed significantly above chance (4.8%). Single asterisk indicates improved classifier performance when combining frequencies or oscillation metrics. Double asterisk indicates further improved performance when combining frequencies and oscillation metrics.

| **rest AB-rest C** |  |  |  |  | *combined across frequencies* |
| --- | --- | --- | --- | --- | --- |
| *phase-sync* | 58.6 | 72.9 | 67.1 | 47.1 | 75.7* |
| *amp-corr* | 32.9 | 62.9 | 62.9 | 50 | 68.6* |
| *power* | 57.1 | 65.7 | 54.3 | 50 | 71.4* |
| *combined across oscillation metrics* | 61.4* | 81.4* | 70.0* | 62.9* | **81.4** |
| **task A-task C** |  |  |  |  |  |
| *phase-sync* | 53.6 | 78.6 | 57.1 | 42.9 | 78.6 |
| *amp-corr* | 35.7 | 71.4 | 39.3 | 57.1 | 64.3 |
| *power* | 50 | 42.9 | 53.6 | 35.7 | 53.6 |
| *combined across oscillation metrics* | 57.1* | 78.6 | 75.0* | 57.1* | **82.1\*\*** |

Our earlier findings highlighted network variability not only across subjects, but also across frequency bands and oscillation metrics *within* an individual. Thus, combining different classifiers that are sensitive to partly non-overlapping information should improve performance. In separate approaches, we fused classifiers across frequency bands, oscillation metrics, or both, separately for rest and task (see Methods). Combining information across frequency bands, we obtained numerically improved performance for all oscillation metrics during rest, with each composite classifier showing greater accuracy than the best performing individual classifier they were based on (Table 1), although similar improvements were not seen for task segment classification. Combining information across oscillation metrics improved classification accuracy for rest segments in all frequency bands, and, for task segments, in three out of four bands. Finally, when we combined all classifiers, performance was further boosted to 81% and 82% for rest and task segments, respectively, correctly identifying the source of 57 out of 70 rest segments and 23 of 28 task segments (binomial tests: both P<10^−16^; permutation tests: both P<0.001). In sum, the improvements observed from these mergers strongly suggest that networks based on different metrics and frequency bands contain unique identifying information.

#### Subject identification

Importantly, successful subject identification does not require correct classification of each individual data segment when multiple segments are available from a subject. Pooling across segments separately for rest and task segments, we correctly identified 13 of the 14 subjects based on rest networks, and 11 of 14 using task networks (binomial tests: both P<10^−13^; permutation: both P<0.001). Task-based classification rates were similar for individual data segments and for subject identity (82% vs. 79%), but the greater number of rest segments available for analysis led to numerically improved subject recognition (93%) relative to segment classification (81%). In a final step, we combined rest and task information, thereby harnessing information from multiple oscillation metrics, frequency bands and behavioral states. Using this approach, we reached perfect accuracy, correctly recognizing 14/14 subjects (binomial test: P<<10^−16^; permutation test: P<0.001). In sum, oscillatory network organization within an individual is sufficiently distinct across frequencies, oscillatory metrics, and behavioral states to differentiate that individual from others, thereby having the potential to serve as a brain-based fingerprint.

#### Influence of frequency bands, oscillation metrics and data segments

We examined the contribution of different network types to our classification results by repeatedly excluding one or more network types from the classifier merger procedure (but retaining both rest and task networks). Removing either all phase-based or all amplitude-based information did not affect performance, but excluding power topographies decreased classification to 86% (12 of 14 subjects). Including only a single oscillation metric (but keeping all frequency bands), subject recognition was 79% for phase-based networks, and 71% for both amplitude-based networks and power topographies. Including only single frequency bands (but retaining all oscillation metrics), we obtained classification rates of 64% for theta, 86% for beta and 79% for gamma. Impressively, including only the alpha band left accuracy at 100%. Using only single oscillation metrics for alpha networks, but still combining rest and task information, resulted in recognition rates of 86% for phase-based, 79% for amplitude-based, and 71% for power-based classifiers. Thus, while alpha activity affords sufficient discriminatory power on its own, these results also demonstrate that alpha networks based on different oscillatory metrics capitalize on different sources of discerning information.

Including all metrics and frequency bands again, we repeated the subject classification procedure using only two rest segments and two task segments (for both training and testing) to match the number of segments available to task classifiers and obtained a success rate of 93% (13 of 14). Next, including all rest and task segments again, we evaluated whether our results might depend on the number of nearest training networks evaluated when labeling a test network. Varying the parameter *k* between 1 and 10 we found that for *k*=1 and *k*=2 one subject was misclassified, but accuracy was stable at 100% for all higher values. Finally, we tested the classifiers trained on Session A/B rest networks on Session C task segments, and vice versa, and combined their votes. This yielded 79% correct subject identification, demonstrating robust network stability across both time and behavioral state.

In summary, these findings demonstrate that better subject recognition is obtained when combining different types of networks, indicating that oscillatory profiles from distinct behavioral states, frequency bands, and oscillation metrics carry unique identifying information.

#### Contribution of individual network components

The analyses described thus far were all based on the full 60-channel EEG data. To examine how many electrodes are required for accurate classification, we varied the number of included elements in each vector used for training and testing classifiers between two and the maximum number available, repeating this process ten times and averaging the results. We first assessed classification rates for individual data segments. Interestingly, the percentage of networks accurately classified by classifiers trained on a single network type reached a plateau quite early on, when approximately 200 out of 1578 (13%) or 100 (6%) connections were included for phase- and amplitude-based networks, respectively (Fig. 6AB). Significantly above chance performance, defined as P<0.05 for one-sample t-tests comparing each sample of ten scores to 4.8%, was achieved with as few as 4.2 ± 1.3 connections for different network types, with alpha amplitude correlation showing significant classification using just two connections (rest: 8.9%, P=0.001; task: 7.9%, P=0.03). For power, performance appeared relatively stable once 20 (33%) electrodes were included (Fig. 6C), but significantly higher than chance performance was observed with only two electrodes for all frequency bands and during both rest and task execution (mean performance: 6.8 ± 1.1%).

**Figure 6.**
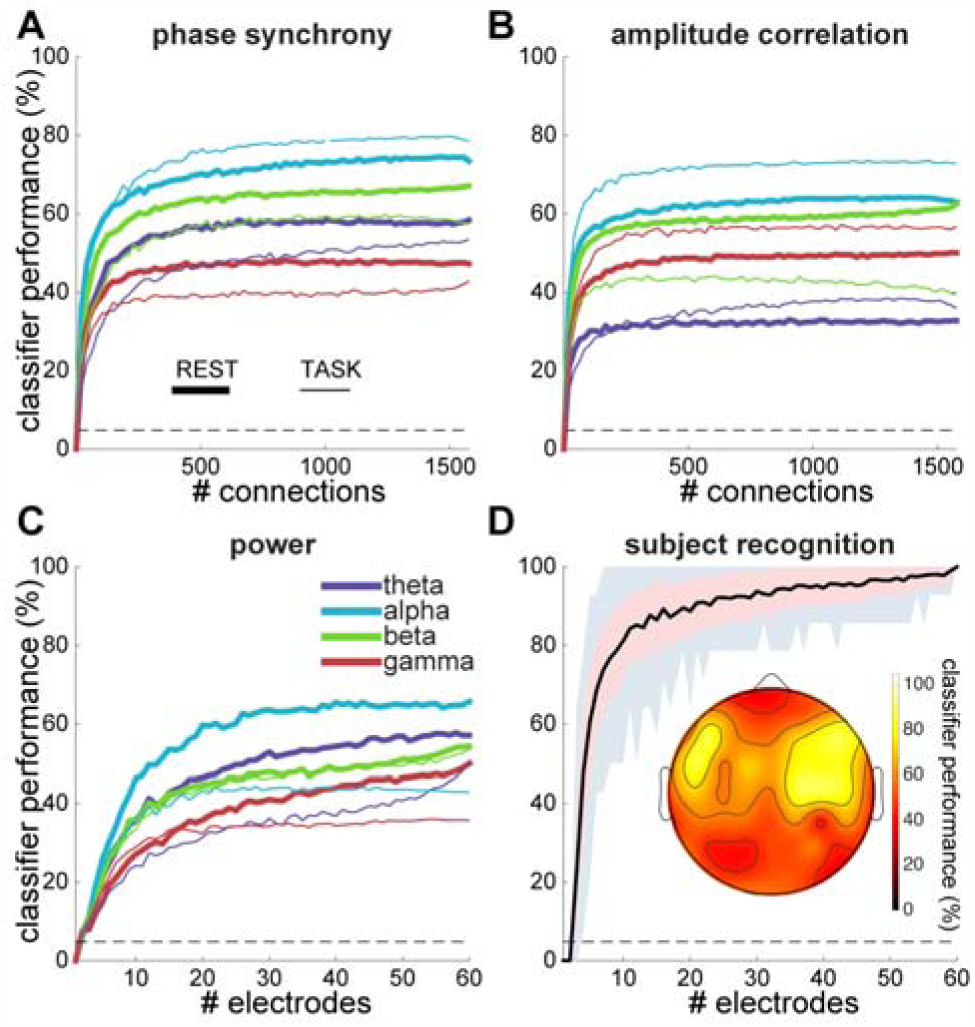
Data segment classification and subject recognition accuracy as a function of number of included connections and electrodes. Percentage of data segments accurately classified as a function of number of included connections for rest and task segments in different frequency bands, for phase synchrony **(A)** and amplitude correlation **(B).** For visualization purposes, A and B data were smoothed with a moving average window of size 11 and down-sampled by a factor 21. Dashed lines indicate chance level performance. **C**: Similar to A and B for power as a function of number of included electrodes. **D:** Subject recognition as a function of electrode array size (electrodes plus connections among them), including all oscillation metrics, frequency bands and behavioral states. Black line indicates average, red shading standard deviation, and grey shading range of minimum and maximum values across 100 iterations. Inset: topographical map displaying subject recognition for searchlight analysis.

Next, we asked how subject recognition rates (i.e., when multiple network types and data segments from the same individual are pooled) depend on these numbers. We varied the number of randomly selected electrodes between two and sixty, and selected all pairwise connections, except neighbors, among them. While the resulting electrode layouts did not reflect conventional montages, they allowed for a simple parametric manipulation of scalp coverage. We repeated this process 100 times for each montage size, training, testing and combining the different classifiers to assess subject identity for every montage size. Results indicated improved performance with larger electrode arrays, with a shape roughly following that of individual classifiers (Fig. 6D). On average, arrays of five, ten and 21 electrodes were sufficient to obtain subject identification rates of 60, 80 and 90%, respectively.

We wondered whether particular clusters of adjacent sensors contributed more to classifier success. To answer this question, we performed a searchlight analysis where, for each electrode, we selected all surrounding electrodes and connections, except those between direct neighbors, within a small radius. Selecting an average of 8 neighbors (range: 6-11) around each searchlight center, we then trained and tested classifiers on these sub-networks, using information from all network types and both behavioral states. Average subject recognition rate across all searchlight centers was 58 ± 14% (range: 36-86%). When we compared searchlight-based recognition rates to recognition scores from random, and therefore generally more distributed, electrode arrays of similar size (9 electrodes), we observed, on average, far superior performance for these distributed networks [79 ± 11%, t(158)=10.2, P<10^−16^]. Topographically, searchlight-based performance was highest at 86% (12 of 14) in two symmetrically lateralized frontocentral clusters, centered on 5 electrodes in total (Fig. 6D, inset). This score was significantly elevated compared to randomly distributed networks of the same size [one-sample t-test: t(99)=6.2,P=10^−8^]. However, topographical maps of individual classifier performance – based on only one network type – were highly variable and generally not suggestive of superior discriminability over particular cortical regions.

In sum, while greater numbers of included connections and electrodes improve subject recognition rates, a remarkable amount of identifying information can be extracted from networks of much smaller size, especially when electrodes are spaced further apart.

## Discussion

In the present work, we offer an extensive analysis of the multivariate network structure of continuous rhythmic brain activity as measured by scalp EEG. Employing a data-driven approach with internal replications, we demonstrate that oscillatory network patterns differ across individuals, behavioral states, frequency bands, and oscillation metrics, suggesting that distinct network types capture separate processing streams operating in parallel. Moreover, while we established that there are clear commonalities in oscillatory templates across subjects, we also observed robust individual differences. Remarkably, individuals exhibited characteristic network profiles that were sufficiently stable over several months to allow successful long-term identification, demonstrating the potential usefulness of topographical patterns of oscillatory activity for biometric purposes.

### Distinct oscillatory profiles based on behavioral state, frequency, and oscillatory feature

Considering these results in more detail, we showed that the network structure of EEG oscillations is reliably distinct for different behavioral conditions, frequency bands, and oscillation metrics. The observation that different behavioral states are associated with distinct spectral profiles is a classic finding [e.g., (Pfurtscheller 1992)] that we also observe (Fig. 2A). Our network similarity approach simply confirms this from a pattern similarity perspective. Variations in the topographical organization of power and connectivity as a function of frequency have also been demonstrated previously (Hipp et al. 2012; Siems et al. 2016). Our similarity findings fit with these observations and extend them by offering a detailed description of how reliably distinct frequency bands differ from each other during rest and task. These observations are further supplemented by investigating several distinct measures of oscillatory brain activity. While the great variety of available spectral and connectivity methods has been widely discussed and simulated (Cohen 2014; Bastos and Schoffelen 2016), direct comparisons of power-, phase-, and amplitude-based activity in human physiological recordings are scarce (Bruns et al. 2000; Arnulfo et al. 2015). We observed reliably distinct oscillatory profiles for large-scale networks based on these metrics, suggesting these measures are indeed sensitive to different kinds of dynamics.

More generally, the notion that several frequency-specific and metric-specific patterns may be derived from the same data supports the idea that EEG activity reflects distinct “layers” of multiplexed activity. Membrane oscillations can coordinate spiking activity in distributed cell assemblies across space and time according to many organizational schemes regarding frequency, phase, and amplitude (Akam and Kullmann 2014; Watrous et al. 2015). To the extent that multiple such oscillatory phenomena contribute to the macroscopic EEG, decomposing EEG signals into their putative components allows identification of parallel modes of neural coding, which, importantly, can serve distinct functional roles (Schyns et al. 2011; Watrous et al. 2013). Although we observed substantial similarity levels even between networks operating in different frequency ranges, or based on distinct oscillatory features, permutation results indicated separable networks well beyond this baseline commonality. Thus, our findings offer physiological support for the notion of multiple, parallel, brain-wide networks.

We observed differentiated networks on two levels. First, at the group-level, our findings indicate that networks of different types (e.g., beta amplitude correlation networks during rest and during task) are statistically separable, yet remarkably consistent across individuals. While most analysis approaches in cognitive neuroscience already assume different brains are sufficiently similar to enable meaningful comparisons between groups (e.g., as a function of behavioral condition or clinical group), there is only limited evidence to suggest such between-subject correspondences hold for macroscopic cortical oscillation structures. Thus, our findings add to a growing literature reporting consistent cortical connectivity patterns across subjects (Chu et al. 2012; Hipp and Siegel 2015; Siems et al. 2016). This finding is perhaps even more notable given the marked between-subject variability we observed. Further supporting these group-level analyses, our demonstration that classifiers trained on oscillatory network structure could successfully differentiate between an out-of-sample subject's rest and task states, and, moreover, could do so for the great majority of frequency bands and oscillation metrics, further underscores the similarity of network patterns across subjects, as well as the difference between these behavioral states. Second, we observed that specific network types were statistically separable even within individuals, highlighting the robustness of our group effects. Furthermore, the presence of several multiplexed oscillatory profiles within an individual importantly supports improved subject recognition by pooling information across frequency bands, oscillatory metrics, and behavioral states, as discussed below.

#### Individual variability and stability of network structure

Alongside the distinctions between network types, and the similarities in oscillatory network organization across subjects, we uncovered substantial individual differences for all network types examined. The distinctiveness of these idiosyncratic oscillatory patterns far surpassed the commonalities shared across subjects. While the suggestion that different brains are organized somewhat differently is not contentious (Mueller et al. 2013; Finn et al. 2015), we demonstrate that EEG-measured network activity is robustly sensitive to such differences. In observing that individual oscillatory profiles persisted, and were in fact identifiable, across months, we conclude that these patterns reflect traits rather than states. It has long been noted that numerous aspects of electrophysiological activity vary among individuals and are, in fact, heritable (Begleiter and Porjesz 2006). Stability of oscillatory parameters across several days to years has been shown for the power spectrum (Salinsky et al. 1991; Kondacs and Szabó 1999), power envelope autocorrelation (Nikulin and Brismar 2004), and graph theoretical characteristics (Deuker et al. 2009; Hardmeier et al. 2014), to name just a few. Oscillatory patterns have also been found to be stable within individuals across frequency bands, brain states, and time (Chu et al. 2012). Our findings build on this body of work and additionally demonstrate oscillatory stability holds across different network types and brain states, both within and across subjects. While it is in principle possible that non-neural factors, such as remaining eye and muscle activity, may have led to our (and others’) findings of long-term stability of EEG signals, we found that pooling information across rest and task networks led to higher classification accuracy compared to using a single behavioral state, suggesting that state-specific neural activity contributed to the observed network stability. Moreover, as we demonstrate in our companion paper (Cox, et al., submitted), individual variability in network organization is robustly related to memory performance, a finding that is difficult to explain unless oscillatory patterns capture cognitively meaningful brain activity.

#### Network-based subject identification

Moving from network stability to identification, many different EEG features have been successfully employed for subject recognition [for an overview, see (Del Pozo-Banos et al. 2014)]. However, only a limited number of studies have addressed the long-term permanence of these effects, relying instead on data from the same acquisition session(s) for both classifier training and testing. This constitutes an important factor because differences in electrode placement across recordings might impact performance substantially. Nonetheless, we achieved highly accurate recognition rates without the use of neuronavigational tools to ensure similar cap positioning across visits, thus indicating a remarkable degree of robustness with respect to precise electrode placement.

While distributed oscillatory patterns have been used for subject identification before (Rocca et al. 2014; Maiorana et al. 2016), our approach substantially increased the interval between recordings. More generally, however, we do not wish to claim that subject identification based on oscillatory patterns leads to the most distinguishable neural fingerprints possible, as we are aware of various other approaches achieving impressive performance rates for EEG- (Näpflin et al. 2007; Lewandowski et al. 2013; Maiorana et al. 2016) or fMRI-based classification (Finn et al. 2015). Rather, by analyzing the different contributions of different frequency bands, oscillation metrics, behavioral states, and the inclusion of specific electrodes/connections, we hope to come to a more thorough understanding of the scope and limits of individual network variability.

Interestingly, the combination of information from distinct network types generally resulted in improved classification rates, indicating that different network types contributed differently. On the whole, power topographies afforded about as much identifying information as connectivity profiles did. This may seem surprising given that power topographies are based on a much smaller number of unique values compared to connectivity vectors. However, connectivity measures are inherently noisier than power estimates, possibly explaining the similar performance levels. While we also noted independent contributions to classification accuracy from different frequencies, alpha activity was sufficient to reach perfect accuracy. This effect is likely at least partly related to the overall prominence of alpha activity, leading to higher signal-to-noise ratios and more accurate (and reproducible) oscillatory estimates. It is unclear whether alpha networks afford superior recognition beyond this consideration. Finally, while task- and rest-based classification rates differed by only a small amount, their combined information enabled us to boost recognition to 100%, indicating the same combinations of frequency bands and oscillation metrics offer some distinct identifying information during different behavioral states.

An open question pertains to the number of unique individuals our approach could conceivably recognize before different subjects’ network structures begin to overlap and reduce performance. In our sample, networks from different individuals show high baseline degrees of similarity, suggesting networks cannot freely occupy arbitrary positions in multidimensional space. Moreover, power and connectivity values between adjacent electrodes and frequency bands are typically correlated, further limiting the number of potential network configurations. Even with these constraints, however, the number of possible network states is enormous. In fact, the dimensionality of this space may be arbitrarily increased by estimating network structure for more fine-grained frequency bins – potentially targeting subject-specific frequencies (Haegens et al. 2014) – by including additional oscillation metrics (e.g., directional connectivity measures), or by expanding the number of cognitive states sampled. Moreover, our results show that substantial reductions of network size still resulted in quite accurate performance, indicating sizeable redundancy across the full network. We speculate that this redundancy is related, in part, to the limited number of individuals we had available, as larger samples might require formerly uninformative connections to assist in differentiating between the increased number of subjects.

#### Limitations and outlook

A potential issue affecting our analyses concerns volume conduction, whereby activity from a single brain source projects to multiple sensors, giving rise to artificially inflated connectivity estimates between nearby electrodes (Palva and Palva 2012). While we used a surface Laplacian filter to reduce volume conduction (Perrin et al. 1989), and removed neighboring channels from the connectivity matrix, we acknowledge these approaches do not completely abolish volume conduction. Yet, if connectivity is driven by volume conduction, power and connectivity should be correlated (Cohen 2014). However, we did not find any evidence for such a relation, neither across individuals, nor across data segments within the same individual (Supplementary Text).

More fundamentally, however, we should stress that the level of analysis in our network approach concerns the degree of similarity between oscillatory activity patterns, which is still meaningful, for our purposes, even if confounded by power. That said, we found that power, phase-, and amplitude-based networks are statistically separable, both within and across individuals, and that connectivity-based networks allowed better subject recognition than power topographies in several instances. Thus, even if we assume a large degree of spurious connectivity, empirical evidence clearly indicates connectivity-based networks to be of practical use beyond that afforded by power topographies. Finally, we note our approach is conceptually similar to multivariate fMRI analyses in which multi-voxel activity patterns (Kriegeskorte 2008), or voxel-voxel correlation patterns are examined (Tambini and Davachi 2013), even though neighboring voxels are typically highly correlated. In sum, while empirical results suggest volume conduction does not pose a major concern for our connectivity results, we emphasize the scope of our network analyses and associated methodological considerations are different from those encountered in conventional EEG analyses.

What, then, is the nature of information indexed by multivariate oscillatory patterns? It is generally accepted that EEG signals reflect the summed activity of synaptic potentials rather than action potentials (Buzsáki et al. 2012; Lopes da Silva 2013), but the precise relation between scalp potentials and underlying brain activity is an active topic of research. Recently, it was found that scalp-recorded EEG activity depends on both the amplitude and synchrony of intracortical sources, but that this relation, in turn, varies with frequency band (Musall et al. 2014). Moreover, oscillations of a particular frequency may be generated by distinct mechanisms in different cortical layers (Bollimunta et al. 2011), and may serve different computational roles in different areas (Supp et al. 2011). Thus, the brain-wide EEG patterns we considered reflect mixtures of many, likely non-linearly, interacting neural processes. Furthermore, in considering only the similarity of oscillatory patterns between conditions, individuals, etc., this approach further abstracts away from particular cortical regions or connections and adopts a more global perspective. Therefore, while our sensor-level network approach does not address anatomically resolved hypotheses as source reconstruction approaches do (Palva et al. 2010; Hipp et al. 2012), its sensitivity to changes in distributed oscillation patterns may offer a complementary perspective on large-scale brain dynamics (Kriegeskorte 2008; Park and Friston 2013; Pessoa 2014; Petersen and Sporns 2015). Indeed, as we report in our companion paper (Cox et al., submitted), distributed patterns of alpha activity relate robustly to individual differences in memory performance, while we do not observe this relation for localized alpha activity. These findings fit with electrophysiological evidence indicating that representational categories may be identified from distributed sensors (Van de Nieuwenhuijzen et al. 2013; Kaneshiro et al. 2015), and fMRI findings that neural regions carrying object category information are quite widespread (Haxby 2001). At the same time, our findings appear quite compatible with previous source-level studies (Hipp et al. 2012; Siems et al. 2016), further lending credence to our approach.

In conclusion, we characterized the variability and stability of large-scale distributed oscillatory networks across individuals, behavioral states, frequency bands, and several theoretically relevant attributes of rhythmic activity. This approach revealed the existence of several canonical, yet distinct, oscillatory profiles, and demonstrated that such patterns constitute neural fingerprints with potential biometric applications. These observations attest both to the human brain's incredibly complex spatiotemporal dynamics and to the wealth of information that can be extracted from EEG signals. We suggest that examining oscillatory activity from a network similarity perspective is a fruitful approach for future studies addressing brain activity and its relation to cognition and behavior.

## Acknowledgements

We thank Alexandra Morgan for technical assistance, and Michael X Cohen and Michael Murphy for valuable comments on earlier versions of this manuscript. This work was supported by grants from The Netherlands Organisation for Scientific Research (NWO) to RC (446-14-009), National Institutes of Health to RS (MH048832), and The Harvard Clinical and Translational Science Center (TR001102).

